# Development of a multi-locus CRISPR gene drive system in budding yeast

**DOI:** 10.1101/391334

**Authors:** Yao Yan, Gregory C. Finnigan

## Abstract

The discovery of CRISPR/Cas gene editing has allowed for major advances in many biomedical disciplines and basic research. One arrangement of this biotechnology, a nuclease-based gene drive, can rapidly deliver a genetic element through a given population and studies in fungi and metazoans have demonstrated the success of such a system. This methodology has the potential to control biological populations and contribute to eradication of insect-borne diseases, agricultural pests, and invasive species. However, there remain challenges in the design, optimization, and implementation of gene drives including concerns regarding biosafety, containment, and control/inhibition. Given the numerous gene drive arrangements possible, there is a growing need for more advanced designs. In this study, we use budding yeast to develop an artificial multi-locus gene drive system. Our minimal setup requires only a single copy of *S. pyogenes* Cas9 and three guide RNAs to propagate three separate gene drives. We demonstrate how this system could be used for targeted allele replacement of native genes and to suppress NHEJ repair systems by modifying DNA Ligase IV. A multi-locus gene drive configuration provides an expanded suite of options for complex attributes including pathway redundancy, combatting evolved resistance, and safeguards for control, inhibition, or reversal of drive action.

## INTRODUCTION

The discovery and implementation of CRISPR/Cas gene editing has revolutionized countless fields and sub-specialties across molecular biology and biotechnology to improve human health, agriculture, ecological control, and beyond. Briefly, alteration of the genetic code is accomplished using (i) a bacterial derived nuclease (typically Cas9 or Cas12a), (ii) a singlestranded fragment of “guide” RNA, and (iii) an optional exogenous repair fragment of DNA (Jinek *et al.* 2012; Jinek *et al.* 2013; Doudna and Charpentier 2014; Zetsche *et al.* 2015). Priming of the nuclease with a pre-programmed guide RNA fragment targets a specific genomic sequence for a double stranded break (DSB). Following DNA cleavage, eukaryotic cells activate repair systems to either fuse broken chromosomal ends together via non-homologous end joining (NHEJ) or, in the presence of donor DNA, introduce exogenous sequence via homologous recombination (HR). Moreover, the CRISPR methodology is not restricted to DSB-induced alteration of the genome—recent efforts have demonstrated that nuclease-dead variants (e.g. dCas9) can serve as delivery systems to modulate transcriptional activity (Qi *et al.* 2013), alter epigenetic landscapes (Thakore *et al.* 2015), or introduce mutational substitutions *sans* any DNA cleavage event (Gaudelli *et al.* 2017).

One powerful biotechnological application of the CRISPR methodology is within a “gene drive” system. The basic design of a homing drive includes the expression constructs for the CRISPR nuclease and the corresponding guide RNA positioned at a desired locus of choice—the mechanism of propagation involves targeting of the homologous chromosome (within a diploid or polyploid organism) at the same genetic position (typically cleaving the wild-type gene). Creation of a DSB followed by HR-based repair (using the gene drive-containing DNA as a donor) causes the entire artificial construct (Cas9, the sgRNA, and any desired “cargo”) to be copied; in this way, a heterozygous cell is automatically converted to the homozygous state. This super-Mendelian genetic arrangement allows for the forced propagation of a genetic element within a population and has the potential to modify entire species on a global scale (BULL AND BARRICK 2017; Godfray *et al.* 2017). Some of the possible benefits of this technology include eradication of invasive species (Esvelt and Gemmell 2017; Prowse *et al.* 2017), agricultural pest management (Courtier-Orgogozo *et al.* 2017), and elimination of insect-borne diseases such as malaria (Godfray *et al.* 2017; Hammond and Galizi 2017; Lambert *et al.* 2018). A number of recent studies have demonstrated the potency and success of CRISPR-based gene drives in fungi (DiCarlo *et al.* 2015; Roggenkamp *et al.* 2017; Roggenkamp *et al.* 2018; Shapiro *et al.* 2018), and metazoans (Gantz *et al.* 2015; Hammond *et al.* 2016; Champer *et al.* 2017; Grunwald *et al.* 2018). While ongoing technical challenges remain in the design, optimization, and field testing of gene drive-harboring organisms, there are also serious biosafety and ethical concerns regarding use of this biotechnology as even current drive systems are expected to be highly invasive within native populations (Noble *et al.* 2018). There is an immediate need for further study (*in silico* and *in vivo*) of gene drive systems that focus on issues of safety (DiCarlo *et al.* 2015; Najjar *et al.* 2017; James *et al.* 2018), control and reversal (Vella *et al.* 2017; Basgall *et al.* 2018), and optimal design (Prowse *et al.* 2017).

There are many types of gene drive designs including “daisy-chain drives,” “underdominance drives,” and “anti-drives,” each with a distinct arrangement of the basic CRISPR components that is predicted to sweep through native populations at varying levels/rates (Godfray *et al.* 2017; Burt and Crisanti 2018; Dhole *et al.* 2018). Moreover, the need for additional drive components (more than one guide RNA construct), genetic safeguards, and built-in redundancy, calls for a new level of complexity within drive architecture. Here, we demonstrate use of *multiple gene drives* across three chromosomal loci within an artificial budding yeast system. Our “minimal” multi-locus gene drive arrangement requires only a single copy of the *S. pyogenes* Cas9 gene (installed at one position), along with three distinct guide RNAs to multiplex the nuclease throughout the genome. We demonstrate that this technique could be used to perform targeted replacement of a native gene (under its endogenous promoter) *in trans* from the Cas9-harboring locus. Finally, reducing or modulating NHEJ by targeting the highly conserved DNA Ligase IV may provide a means to further bias HR-dependent repair and action of gene drives across eukaryotic systems. Our method includes multiple layers of genetic safeguards as well as recommendations for future designs of multi-locus drive systems.

## RESULTS

### Rationale and design of a multi-locus CRISPR gene drive

To date, a number of studies in fungi, insects, and now vertebrates, have demonstrated that CRISPR-based gene drive systems are effective in both single-celled and multicellular eukaryotes (DiCarlo *et al.* 2015; Gantz *et al.* 2015; Hammond *et al.* 2016; Champer *et al.* 2017; Grunwald *et al.* 2018; Roggenkamp *et al.* 2018; Shapiro *et al.* 2018). One of the benefits of homing systems is the ability to install additional genetic “cargo” proximal to the gene drive (consisting of a nuclease gene and an expression cassette for the guide RNA). Current strategies use the gene drive cassette itself to delete and replace an endogenous gene, and/or include exogenous material as a desired cargo. However, there are a number of limitations to the use of a single locus harboring the entirety of the gene drive. First, addition of entire genetic pathways or large numbers of gene expression systems may be less efficient at HR-based copying of the drive. Second, introduction of additional endogenous gene(s) or modified alleles may require the native promoter system and/or epigenetic landscape to provide accurate and timely expression—this would not be possible at a single generic drive-containing locus. Third, given the observation of both natural (e.g. single nucleotide polymorphisms) and evolved resistance to gene drives (indels resulting from NHEJ) within insect populations (Champer *et al.* 2017; Drury *et al.* 2017; Hammond *et al.* 2017; Unckless *et al.* 2017; Buchman *et al.* 2018; KaramiNejadRanjbar *et al.* 2018), mechanisms for fortifying drive systems are still being elucidated. The proposal to increase the number of targeted double stranded breaks (and corresponding sgRNAs) to the single nuclease of choice (e.g. *S. pyogenes* Cas9) would greatly aid in combatting resistance (Bull and Malik 2017; Marshall *et al.* 2017; Noble *et al.* 2017). However, an independent means to both minimize or escape resistance *and* ensure the intended biological outcome (deletion of the intended gene or introduction of the exogenous cargo) would involve a redundant delivery system. In this way, multiple gene drives (with multiple guide RNAs) within the same organism could target independent genetic loci either from the same, distinct, or parallel genetic pathways to achieve the desired outcome(s).

We envisioned two general strategies for the development of a gene drive system across distinct chromosomal positions: (i) each multi-locus “Complete” Gene Drive (CGD) would contain both a nuclease and corresponding guide RNA or (ii) a multi-locus “Minimal” Gene Drive (MGD) would include a nuclease and sgRNA, and all other genetic loci would *only* contain additional guide RNA cassettes (Fig. 1A, left). We chose to focus on the latter strategy for a number of reasons, but we recognize that both would have distinct challenges and advantages. For one, a possible technical hurdle to development of a modified organism with multiple CGDs would be the generation of distinct “large” expression system consisting of the entire nuclease gene, flanking UTR, the guide expression cassette(s), and any optional cargo compared to the MGD which removes the bulk of the drive system (nuclease expression) at additional loci.

**Figure 1.**
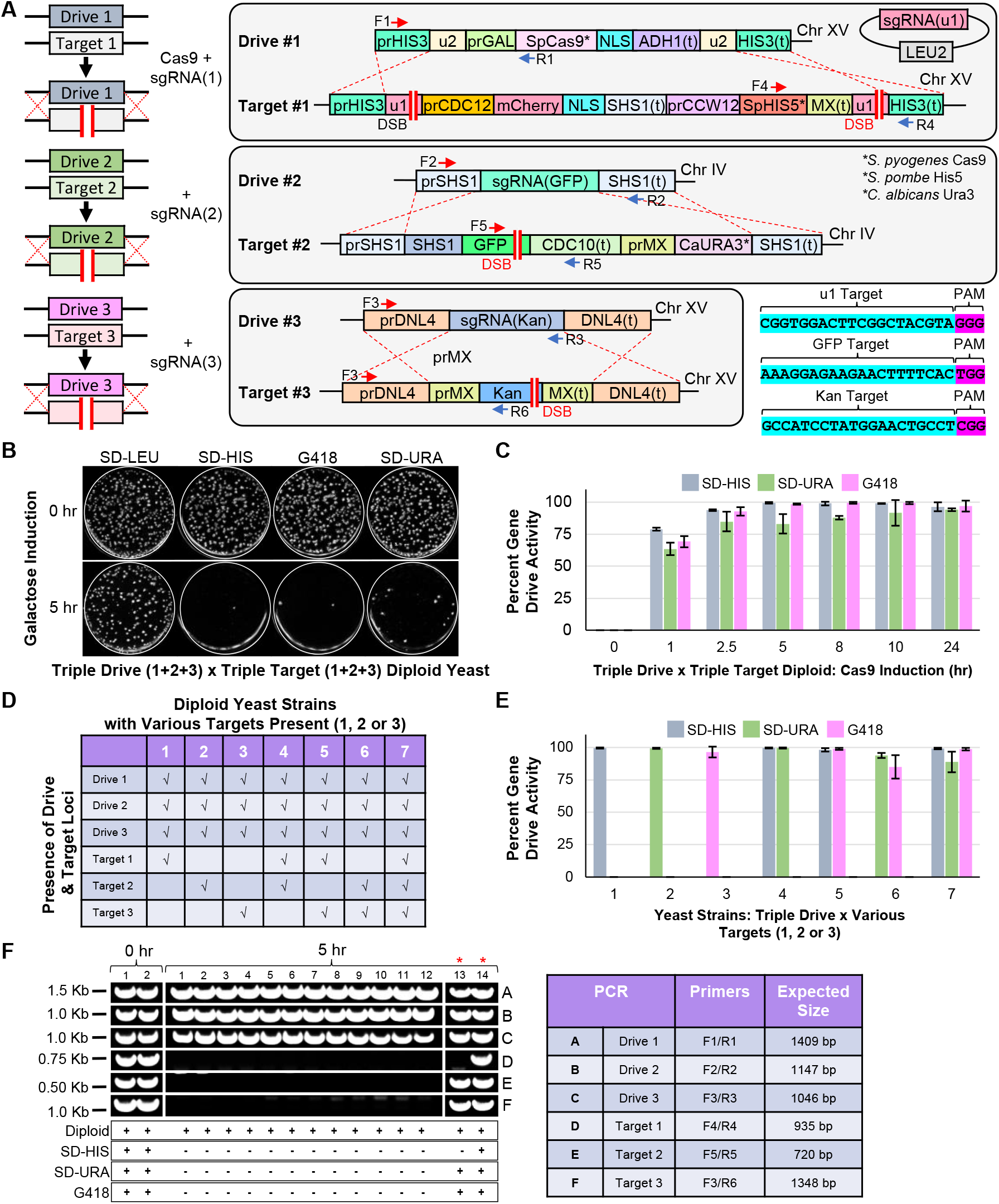
Design of a CRISPR/Cas9-based gene drive system in *S. cerevisiae* across three loci. (A) *Left*, An artificial gene drive was installed at three separate loci in haploid yeast. Each drive system (Drive 1-3) contained a guide RNA cassette targeting an artificial target (Target 1-3) at the same locus. Only Drive 1 contained the cassette for *S. pyogenes* Cas9. *Right*, Artificial (u1) and (u2) sites (Finnigan and Thorner 2016) were used flanking the gene drive at the *HIS3* locus (Chromosome XV) and the *S.p.HIS5* selectable marker. The *SHS1* locus (Chromosome IV) included a C-terminal GFP fusion and the *C.a.URA3* marker. *DNL4* (Chromosome XV) was deleted with the Kan^R^ cassette. All sgRNAs were targeted to non-native sites (sequences shown). The sgRNA(u1) cassette was included on a high-copy plasmid (*LEU2* marker). *S.p.* Cas9 was under control of the inducible *GAL1/10* promoter. (B) Haploid yeast harboring the triple drive (GFY-3675) were mated to yeast of the opposite mating type containing the triple target (GFY-3596) to form diploids. Cas9 expression was induced by growth in galactose for either 0 or 5 hr. Yeast were diluted to 100-500 cells per plate, grown for 2 days, and transferred to SD-LEU, SD-HIS, SD-URA, and G418 plates. (C) A time course of galactose activation using the [GFY-3675 x GFY-3596] diploid in triplicate. Error, SD. (D) Seven haploid yeast strains (GFY-3206, 3593, 3264b, 3578, 3594, 3623, and 3596) were constructed and tested as in (B) against the triple drive strain (GFY-3675). (E) Each of the diploids created from (D) were cultured for 5 hr and quantified for drive success. Error, SD. (F) Clonal isolates were obtained from diploids generated in (B) at either 0 hr (2 isolates) or 5 hr activation of Cas9 (14 isolates). All yeast were confirmed as diploids and assayed on each media type (*below).* Diagnostic PCRs were performed on genomic DNA to detect the presence (or absence) of each drive and target locus; oligonucleotide (Supplementary Table 5) positions can be found in (A) and the expected PCR fragment sizes are illustrated (right). Two isolates (13,14) were chosen for their incomplete growth profile (red asterisks).

Along these lines, the issue of appropriate expression of each of the nuclease gene(s) (whether identical or distinct) would need to be addressed using identical or modified promoter elements; this issue does not exist for a MGD with only a single copy of Cas9. Second, the issue of biosecurity and safeguarding against accidental or malicious release was taken into consideration. Given that a MGD would only harbor one copy of the nuclease, it would provide far less hurdles to counter and inactivate—either through use of an anti-drive system (DiCarlo *et al.* 2015; Vella *et al.* 2017), by induced self-excision (Roggenkamp *et al.* 2018), or by removal of the Cas9-containing drive guide RNA (Roggenkamp *et al.* 2018). Therefore, we have chosen to focus our study on design and testing of a three-locus MGD in budding yeast using the *S. pyogenes* Cas9 nuclease.

### An efficient triple gene drive system functions independently at each locus

Our novel system includes the most potent genetic safeguard known to date used within a gene drive: artificial and non-native sequences used as targets. In this way, we have not only generated a haploid yeast strain harboring the MGD system at three genetic loci (*HIS3, SHS1*, and *DNL4*), but have also created a corresponding haploid strain with three distinct artificial *targets* at the same three loci (Fig. 1A, *right*). The “primary” drive at the *HIS3* locus includes (i) Cas9 under an inducible promoter (*GAL1/10*) commonly used for overexpression, (ii) flanking (u2) artificial sequences to be used for self-excision as a safeguard, and (iii) the absence of any selectable marker. The corresponding guide RNA cassette was installed on a high-copy plasmid for security reasons, but could have also been integrated proximal to the drive itself. The “secondary” and “tertiary” drive systems (*SHS1* and *DNL4*) are both non-essential genes and contain the minimum required components in the MGD design; in both cases, the native gene was deleted and fully replaced by the guide expression cassette (455 bp, although this could be reduced further) with no selectable marker. Construction of this complex haploid yeast strain used a combination of traditional HR-based integrations (with selectable markers), universal Cas9-targeting systems (CRISPR-UnLOCK) (Roggenkamp *et al.* 2017), and novel “self-editing” integration events (Supplementary Fig. 1). To test the efficacy of the MGD, a three-locus “target” strain was generated: the *HIS3* locus was flanked by two (u1) sequences and included the *S.p.HIS5* selectable marker, the *SHS1* gene was fused with GFP and contained the *C.a. URA3* marker, and finally, *DNL4* locus was deleted and replaced with the Kan^R^ drug cassette (Fig. 1A, *right*).

The triple MGD strain was first mated with the triple target strain to form a diploid, and Cas9 was activated by culturing in medium containing galactose (Fig. 1B). In the absence of nuclease expression (Fig. 1B, *top*), all yeast colonies contained the (u1) guide plasmid (*LEU2*), and three selectable markers (*HIS5, URA3*, and Kan^R^—providing resistance to G418). However, following a 5 hr incubation in galactose, >95% of all colonies were sensitive to all three growth conditions indicating a loss of all three selectable markers and replacement via the MGD (Fig. 1B, *bottom*). A time course of galactose induction illustrated highly efficient drive activity for all three loci by five hours; we noticed a slight lag in efficiency for the loss of the *URA3* marker (*SHS1*) locus until the 24 hr mark (Fig. 1C). This observation may be due to the *HIS3* and *DNL4* loci both being present on chromosome XV whereas *SHS1* was located on chromosome IV. Alternatively, differences in available guide RNAs (plasmid-borne versus integrated) or local epigenetic effects could cause this slight reduction in editing. Next, to ensure that action of the MGD at each locus was not dependent on the presence or absence of one or more of the intended targets (simulating “resistance” at one or more loci), we retested the triple drive strain against six additional strains, each lacking one or two of the proper targets and instead, contained the native yeast sequence: *his3A1, SHS1*, or *DNL4* (Fig 1D). We obtained similar results for each combination as the triple MGD strain (#7) indicating that each gene drive functioned independent of the presence of additional target(s) (Fig. 1E). We also observed that drive success at the *SHS1* locus slightly increased when fewer targets were presented. Finally, to ensure that the loss of the selectable marker was coupled to replacement of the target locus by the drive locus, we isolated clonal yeast from the MGD triple cross (Fig. 1B) and confirmed both the growth profile and ploidy status of random samples (Fig. 1F, *bottom*). Diagnostic PCRs were performed on all six distinct loci to assay for the presence or absence of each engineered drive and target (Fig. 1F, Supplementary Fig. 3). Oligonucleotides (Supplementary Table 5) unique to specific drive/target elements were chosen; prior to Cas9 activation (0 hr), diploids contain all six distinct loci (two isolates). However, following activation of the nuclease, diploids maintained all three drive loci (PCRs A,B, and C), but lost all three target loci (PCRs D, E, and F) (twelve independent isolates).

We recognized that following the 5 hr drive activation, a small number of yeast colonies (<5% in most cases) still contained one or more selectable marker(s). We reasoned that these rare colonies likely arose from either complete or partial failure of the gene drive system for various possible reasons (poor expression, loss of guide RNA plasmid, NHEJ, alterations in ploidy, etc.). Therefore, we isolated and tested additional clones that displayed incomplete growth profiles across the three selection plates (isolates 13,14) (Fig. 1F). One isolate (13) had lost the (u1)-flanked target at the *HIS3* locus yet still contained the *SHS1* and *DNL4* markers. The second isolate (14) appeared to have lost the LEU2-based plasmid and all three target loci were still present (Fig. 1F). Following transformation with the (u1) guide vector, we examined a second round of drive activation from these two isolates and obtained a similar growth profile with a loss of the remaining loci indicating that at least some of the “failed” drive occurrences resulted from improper activation and/or targeting (Supplementary Fig. 4). Of note, our gene drive system was activated in the absence of any selection—diploids were grown in rich medium containing galactose, and grown on SD-LEU plates prior to testing of the drive status on various medium. In this way, the action of the gene drive was performed in the absence of any selection or challenge.

### DNA Ligase IV as a target for gene drives

Our choice of the yeast *DNL4* gene as one of the MGD targets was intended to highlight the ability of a drive itself to modify or eliminate non-homologous end joining (NHEJ)—the DNA repair process that directly counteracts the action of gene drives. Following DSB formation by Cas9, the function of the homing drive requires repair of the broken chromosome via homology directed repair using the homologous chromosome (and drive itself) as the source of the donor DNA. However, should NHEJ repair systems ligate the broken chromosome ends prior to HR-based copying, the drive will fail to copy; in fact, imprecise repair by NHEJ may even generate alleles of the target that would be resistant to further rounds of editing. Therefore, this competing DNA repair system remains one major technical hurdle to optimal gene drive design in higher eukaryotes. Of note, interest in modulating, tuning, or inhibiting NHEJ-based repair pathways is not unique to CRISPR gene drives as this mode of repair still competes with the introduction of exogenous DNA via HR (Chu *et al.* 2015; Maruyama *et al.* 2015; Robert *et al.* 2015; Vartak and Raghavan 2015; Schwartz *et al.* 2017; Canny *et al.* 2018).

The NHEJ pathway is highly conserved from yeast to humans and functions to directly fuse exposed DNA ends (Lieber 2010; Chiruvella *et al.* 2013b). DNA Ligase IV (Dnl4 in yeast, Lig4 in humans) is required for the final step of DNA ligation along with other conserved binding partners (Ellenberger and Tomkinson 2008). We examined the genomes of other fungi and metazoans using the yeast Dnl4 protein sequence as a query and a phylogenetic history of this enzyme illustrated the evolution of this enzyme through deep time (Fig. 2A). Note, the branching of *Z. nevadensis* (termite) was poorly supported and has been previously shown to be included within the *Insecta* class (Misof *et al.* 2014). The DNA Ligase IV enzyme is divided into multiple subdomains including DNA binding, adenylation, oligonucleotide binding, and a C-terminal BRCT domain that interacts with binding partner Lif1 (XRCC4 in human). A previous study identified a number of mutational substitutions within the C-terminus of yeast Dnl4 that resulted in a *partial* loss of function of NHEJ (Chiruvella *et al.* 2014). Examination of protein sequence alignments between yeast, mosquito, and human DNA Ligase IV C-terminal domains revealed only a minor conservation of sequence identity (Fig. 2B). However, several of the identified yeast residues were conserved by either insects and/or humans (yeast T744, D800, G868, and G869).

**Figure 2.**
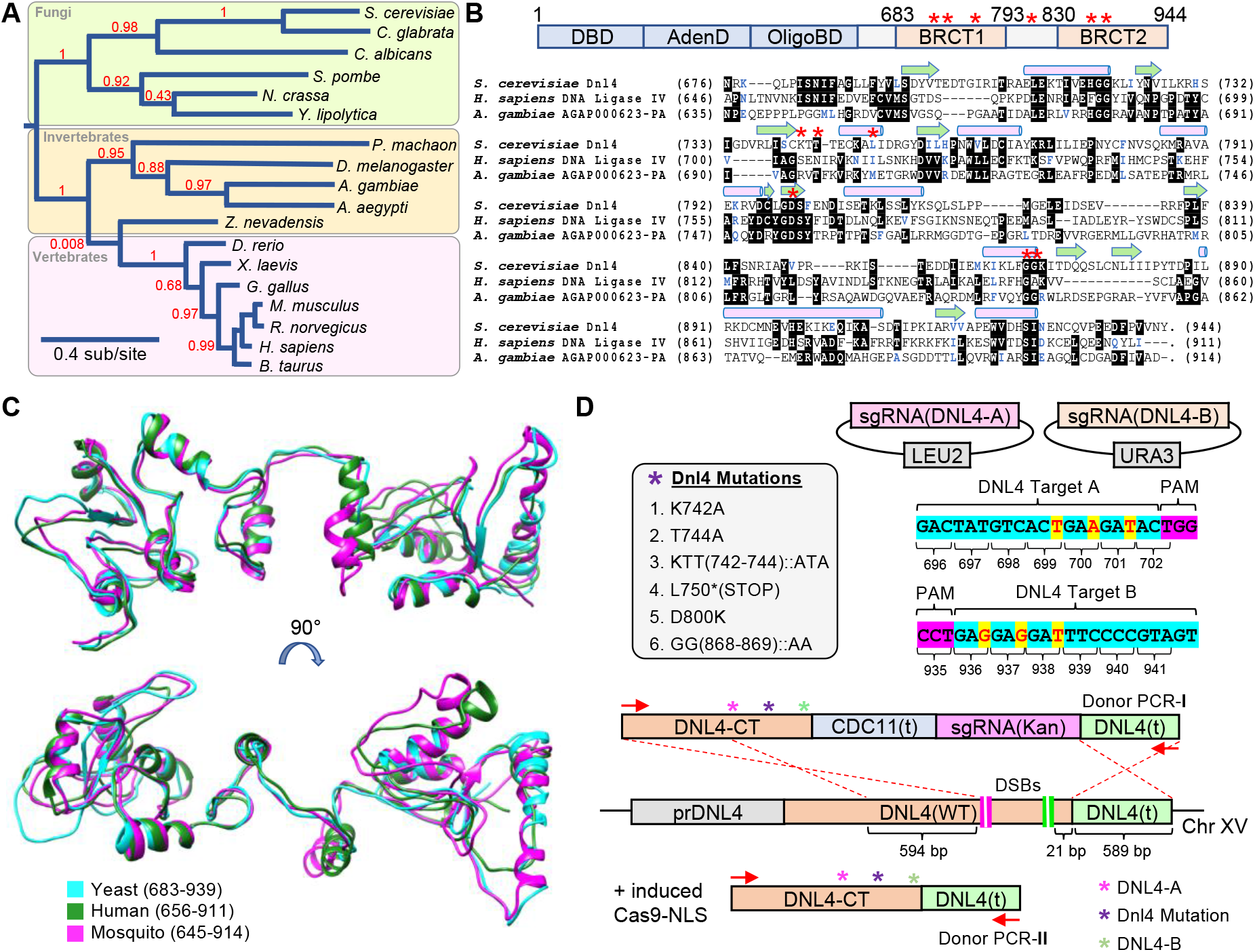
DNA Ligase IV, critical for NHEJ and conserved across eukaryotes, provides a unique candidate for gene drives. (A) Phylogenic analysis of Ligase IV candidates (Supplementary Table 3) across fungi and metazoans by Phylogeny.fr (Dereeper *et al.* 2008; Dereeper *et al.* 2010). Branch lengths correspond to the number of substitutions per site and the confidence of most branches is illustrated as a decimal (red text). (B) *Top*, Illustration of the domain structure of yeast Dnl4. The catalytic N-terminal portion includes a DNA binding domain, adenylation domain, and oligonucleotide domain (blue). The C-terminal portion includes tandem BRCA1 C-Terminal domains (BRCT). *Bottom*, A multiple sequence alignment was performed using Clustal Omega (LI *et al.* 2015) of the yeast, mosquito, and human Ligase IV protein C-termini. Identical residues are shown against a black background and similar residues are colored in blue. Secondary structures (pink cylinder, α-helix; green arrow, β-strand) for the yeast Dnl4 C-terminal as determined by the crystal structure are illustrated (DORE *et al.* 2006). The position of six alleles (K742, T744, L750, D800, G868, and G869) are also illustrated (red asterisk) that were identified from a previous study (Chiruvella *et al.* 2014). (C) The protein sequences of the *A. gambiae* (645-914) and *H. sapiens* (656-911) Ligase IV were modeled against the crystal structure of the *S. cerevisiae* (683-939) Dnl4 (PDB:1Z56) using I-TASSER (ROY *et al.* 2010) (Supplementary Table 4) and illustrated using Chimera (Pettersen *et al.* 2004). (D) Cas9-based genomic integration methodology for introduction of mutational substitutions to the native *DNL4* locus in yeast. Two sgRNA-expressing cassettes were cloned onto high copy plasmids (marked with *LEU2* and *URA3*) to induce two DSBs within the C-terminus of *DNL4.* Silent substitutions were generated within the intended repair DNA to prevent re-targeting of Cas9 (silent alterations in yellow). Two repair strategies were used to include either a non-native terminator coupled with a sgRNA(Kan) cassette, or the native *DNL4* terminator; the included amount of homology (bp) is illustrated.

Using the crystal structure of the C-terminus of yeast Dnl4 as a template, we generated models (I-TASSER) for the corresponding domains of mosquito and human Lig4—both displayed a much higher conservation of structure as opposed to primary sequence (Fig. 2C, Supplementary Table 4). The N-terminal region also displayed strong structural homology using the human Lig4 crystal structure as a template (Supplementary Fig. 5).

While total loss of NHEJ (e.g. *dnl4*Δ) is tolerated in yeast, it is unclear whether a DNA Ligase IV null allele would be viable in higher eukaryotes. Along these lines, truncations or mutations of Lig4 in humans can lead to the rare DNA Ligase IV syndrome (Chistiakov 2010; Altmann and Gennery 2016). However, given that reduction in transcript or replacement by a partially functioning allele could reduce, but not eliminate NHEJ repair, it could be utilized in other systems to maximize gene drive efficiency, even at the (potential) expense of overall fitness. Therefore, we utilized a “self-editing” methodology to integrate six *dnl4* alleles—five partial loss of function substitutions, one truncation, and a WT control (Fig. 2D). In a strain harboring integrated Cas9 at the *HIS3* locus (Roggenkamp *et al.* 2018), we introduced two DSBs within the C-terminus of native *DNL4* and integrated two different constructs: (i) a modified *dnl4* allele with a sgRNA(Kan) cassette and (ii) a modified *dnl4* locus using the native terminator sequence. Both Cas9 target sites were also mutated within the repair (donor) DNA to prevent subsequent rounds of unintended editing.

We utilized these eight haploid strains to quantify the level of NHEJ repair (Fig. 3). Our system of DSB formation followed by DNA repair utilized the dual programmed (u2) sites flanking the Cas9 expression cassette (Fig. 3A). With only a single guide construct, Cas9 would be multiplexed to both identical sites, causing complete excision of the nuclease gene and Kan^R^ marker. Following transformation of the sgRNA(u2) plasmid, yeast were analyzed for the number of surviving colonies on SD-LEU medium (Fig. 3B). Editing by Cas9 at both (u2) sites followed by precise DNA ligation of the broken ends would generate a “new” (u2) site, and would be subject to a second round of Cas9-dependent cleavage—continual DSB formation followed by exacting repair causes inviability in yeast (Roggenkamp *et al.* 2018).

**Figure 3.**
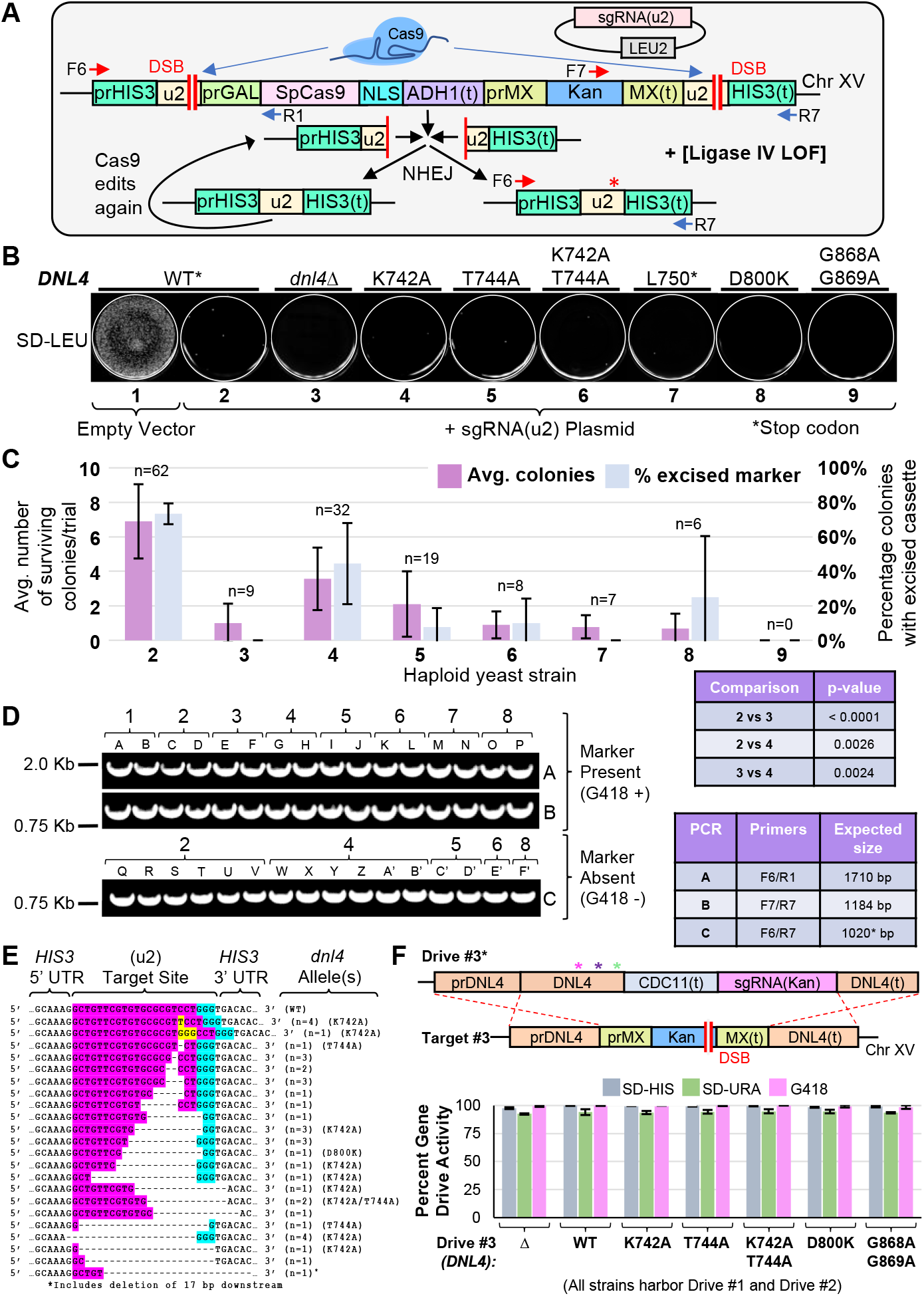
Partial loss of function alleles of yeast DNA Ligase IV reduce NHEJ. (A) Design of a self-excising Cas9-based assay for NHEJ. Strain GFY-2383 included an inducible Cas9 cassette paired with the Kan^R^ marker. Transformation of the sgRNA(u2) plasmid would result in multiplexing to two flanking (u2) sites. Repair via NHEJ would result in the formation of the original (u2) site, and would be subject to further rounds of editing; introduction of an indel (red asterisk) would cause destruction of the Cas9 target site. (B) Strains GFY-3850 through GFY-3856 and GFY-3864 (Supplementary Table 1, *Conditions* 1-9) were transformed with the sgRNA(u2) plasmid (pGF-V809) or empty vector control (pRS425) and plated onto SD-LEU for three days. The *DNL4* (WT) gene contained six silent substitutions (asterisk). (C) The average number of surviving colonies was quantified for all trials—labeled as in (B); the number of colonies (n) obtained across all experiments is displayed. Error, SD. For the pRS425 vector, 2897 +/− 357 colonies were obtained. The percentage of isolates that excised the Cas9 cassette at the *HIS3* locus (by sensitivity to G418) is displayed. Error, SD. Statistical analyses of strain comparisons (colonies per trial) were performed using an unpaired t-test. (D) Diagnostic PCRs were performed on chromosomal DNA from clonal isolates from (B) to illustrate presence (2 isolates each) or loss (between 2-6 shown) of the Cas9 cassette. *Conditions* 2-8 correspond to those found in (B). Oligonucleotides (Supplementary Table 5) used can be found in (A) and the expected PCR sizes are illustrated (right). (E) DNA sequencing of the *HIS3* locus following NHEJ (on isolates sensitive to G418). For each insertion or deletion, the number of identical clones is displayed. All sequences were obtained from WT *DNL4* yeast except unless otherwise noted. Target, pink. PAM, blue. Insertions, yellow. (F) Triple-drive containing strains were constructed with a modified *DNL4* locus coupled with the sgRNA(Kan) cassette. Haploid drive strains (GFY-3675, 3865-3867, 3871, 3872, and 3875) containing the sgRNA(u1) plasmid were mated with GFY-3596, diploids selected, and gene drives activated as in Fig. 1B. The percentage of colonies sensitive to each condition represented gene drive activity. Error, SD.

However, introduction of an insertion, deletion, or substitution within the target sequence would render the site immune from subsequent rounds of editing. Furthermore, loss of the Kan^R^ marker provided a growth phenotype associated with targeting of the (u2) sites and excision of the entire cassette at the *HIS3* locus. Both the total number of surviving colonies as well as the percentage of isolates with an excised marker were quantified in triplicate (Fig. 3C). In our assay, the presence of WT *DNL4* allowed for approximately 7 colonies/experimental trial, whereas *dnl4*Δ yeast resulted in 0-1 colonies on average. Importantly, of the WT *DNL4* isolates, 73% had properly excised the entire cassette whereas this was found to be 0% for *dnl4*Δ yeast across numerous independent trials (Fig. 3C). The partial loss of function *dnl4* alleles provided a range of NHEJ efficiencies: the K742A mutant averaged 4 colonies/trial with an excision rate of nearly 50% and other substitutions displayed excision rates of between 0-25%. As expected, the C-terminal *dnl4* truncation at L750 phenocopied the null allele. Diagnostic PCRs confirmed the presence or absence of the Cas9-Kan^R^ expression cassette for clonal isolates from each of the aforementioned haploid strains tested (Fig. 3D). For strains that had undergone editing and marker excision, the *HIS3* locus was amplified and sequenced; NHEJ followed by imprecise ligation introduced either insertions or deletions at the site of Cas9 cleavage 3 bp upstream of the 5’ end of the PAM sequence (Fig. 3E). Finally, each of the *dnl4* alleles was tested within our MGD system as a native cargo-based delivery system (Fig. 3F, top). Given that our artificial *DNL4* target was the *dnl4*ΔKan^R^ null allele, we recognize that in the context of a [gene drive x WT] diploid genome, further modifications would be required to bias the HR-based repair of the intended *dnl4* allele. This could include recoding (silent substitutions) of the *DNL4* C-terminal domain sequence to prevent promiscuous cross-over downstream of the intended mutation(s). Following expression of Cas9 and activation of the MGD, the growth profiles of 7 diploid strains were assessed in triplicate and demonstrated efficient drive activity at all three loci (Fig. 3F, *bottom*). Moreover, PCRs from clonal isolates confirmed the presence or absence of each drive and target locus (Supplementary Fig. 6). These data demonstrate that the MGD strategy can be used as a knock-out or allele replacement strategy (at a native locus) with only minimal added sequence (782 bp).

## DISCUSSION

In this study, we have developed a multi-locus CRISPR gene drive with a minimal design (MGD) that allows for multiplexing of Cas9 *in trans* across three distinct chromosomal locations (Fig. 4). An alternative strategy could also be employed to create more than one gene drive system within a single genome—a CGD where each locus of interest contains the full complement of genetic information (nuclease, UTR, sgRNA, and optional cargo). In this way, each drive would be completely independent from all other drive(s). While this design clearly provides a maximum level of potential redundancy, there are other technical and safety issues inherent to this multi-nuclease arrangement. For one, countering or inhibiting a CGD with more than one active nuclease would require more sophisticated anti-drive systems, the discovery of additional anti-CRISPR proteins, or complex regulatory systems to ensure inactivation or destruction of each drive. In contrast, our minimal GD design can be inhibited by the AcrIIA2/A4 proteins (Basgall *et al.* 2018), self-excised by our flanking (u2) sites (Roggenkamp *et al.* 2018), or targeted by an anti-drive system no different than a traditional single-locus gene drive. We argue that this type of design provides a higher level of biosecurity and can still accomplish the same task as n-number of “full” gene drives. Moreover, the issue of tightly regulated control of the nuclease transcript may pose additional challenges if the same promoter elements are positioned across multiple chromosomes and epigenetic landscapes in the CGD design.

**Figure 4.**
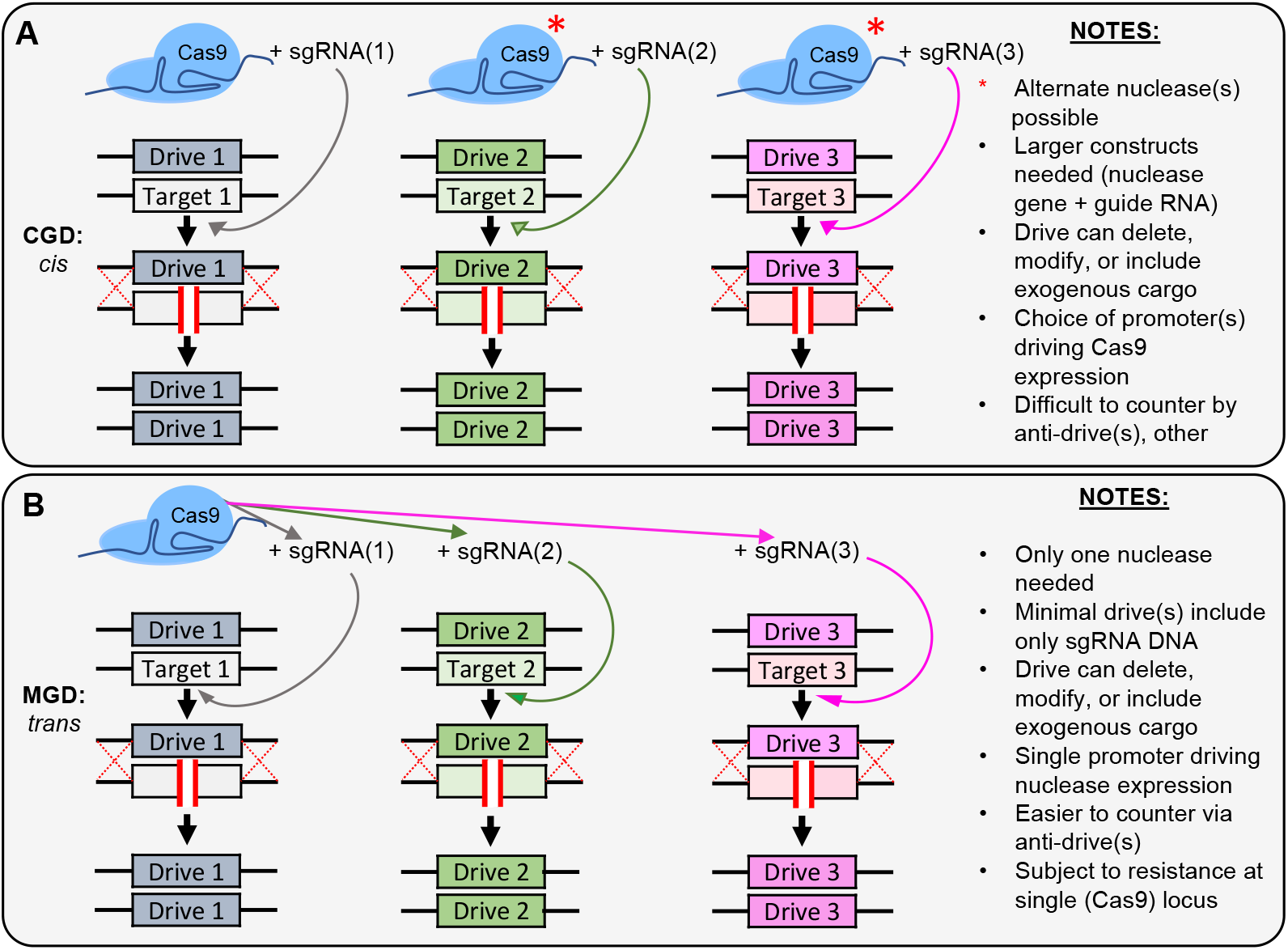
Models for multi-locus CRISPR gene drive systems. (A) A proposed gene drive arrangement *in cis.* Each locus to be modified contains a “complete” system (drive and guide RNA cassette). These may be identical nuclease genes, altered variants, or sourced from separate species (e.g. Cas9 versus Cas12a). The action of each drive is fully independent from other drive-containing loci. (B) A single nuclease functions *in trans* across multiple loci with separate guide RNAs. This “minimal” design allows for greater safety and security (easily countered by a single anti-drive system or other means) but may be more susceptible to resistance at the primary (Cas9-harboring) locus.

One potential issue facing our MGD design (or *any* gene drive design, for that matter) is that of natural or evolved resistance to the action of the drive. Since our multi-locus arrangement includes only a single nuclease gene powering all drives, any resistance or escaped action to the Cas9-containing locus would render all three gene drives inactivated in subsequent generations. However, evidence now exists both *in silico* (Marshall *et al.* 2017; Noble *et al.* 2017; Prowse *et al.* 2017) and *in vivo* (Champer *et al.* 2018) that the addition of multiple guide RNAs (to the same genetic target) can reduce (or potentially *eliminate*) resistance to the gene drive. Using current estimations for a given target, five separate guide RNAs may provide a sufficiently rare or improbable event requiring mismatch or mutation to occur at all five target DNA sites (Marshall *et al.* 2017; Noble *et al.* 2017). This would provide greater than 99% confidence in eliminating an *A. gambiae* population on a continent-wide scale (Marshall *et al.* 2017). Therefore, our recommendation would be to greatly bias multiplexing (via multiple guide RNAs) to the gene drive locus harboring the sole copy of Cas9 in the MGD design (in our system, Cas9 creates two DSBs flanking the entire locus). To ensure even higher fidelity of the nuclease and to combat resistance, one could combine the two strategies (CGD and MGD) to have a secondary copy of the nuclease (of the same variant or a different species) positioned at a second locus—additional multiplexing across numerous other loci could include the minimal design (guide RNA cassette only). Finally, while our MGD methodology includes a gene drive consisting of only 455 nucleotides (sgRNA expression cassette), this could be reduced even further to be only a few bases or the absence of any base pairs. Additional sgRNA cassettes could be installed at one or more loci to allow for targeting of chromosomal positions where the “drive” is nothing more than a single base substitution or deletion. Provided few bases separate the DSB site and the intended mutation(s), HR-based repair would allow for propagation of the few bases no different than a “full” gene drive (consisting of many thousands or tens of thousands of bases) into the homozygous condition. The only requirement would involve the sgRNA expression cassette(s) to also be installed within a drive-containing locus *in trans.*

Finally, we chose one of the chromosomal targets within our minimal gene drive system to include both deletions, truncations, and substitution alleles of *DNL4*—one of the essential components of the NHEJ repair pathway. While loss of DNA Ligase IV is non-lethal in yeast (Wilson *et al.* 1997) and flies (Gorski *et al.* 2003), it is embryonic lethal in mouse (Barnes *et al.* 1998) and not tolerated in mosquito (Basu *et al.* 2015). However, suppression or inhibition of this enzyme or the NHEJ repair pathway has been shown to increase rates of recombination and genomic integration of exogenous DNA using CRISPR systems *in vivo* (Basu *et al.* 2015; Chu *et al.* 2015; Maruyama *et al.* 2015; Robert *et al.* 2015; Vartak and Raghavan 2015; Cen *et al.* 2017; Canny *et al.* 2018). We demonstrate that this critical factor could be an additional target for a multi-locus gene drive system—suppression of NHEJ, whether by mutated alleles, regulation of transcript, or direct inhibition of the enzyme—would aid in successful HR-based copying of the drive and further reduce the possibility for drive resistance, especially when coupled with multiple guide RNAs. Numerous strategies might be employed to accomplish targeted suppression of NHEJ including testing of additional Ligase IV loss of function alleles that may be widely conserved across eukaryotes; our study focused on the C-terminal BRCT-domain containing portion of Dnl4, but other substitutions have been also been characterized within the N-terminal catalytic domain (Chiruvella *et al.* 2013a).

A multi-locus CRISPR gene drive system should help advance current designs and provide additional options for (i) biosecurity, (ii) drive redundancy, (iii) combatting of evolved resistance, (iv) native gene replacement, (v) multiple gene cargo/genetic pathway delivery, (vi) suppression of NHEJ or activation of HR-promoting repair pathways, and (vii) multiple phenotypic outcomes. Advanced drive arrangements (Dhole *et al.* 2018) could accomplish multiple outcomes within a single-genome system—the additional of exogenous cargo could also be paired with (native) allele introduction *and* modulating of organism fitness by perturbing numerous other genetic pathways in a single step. As the design and application of CRISPR gene drives continues to advance, we continues to stress the need for multiple levels of control, tunability, inhibition, and drive reversal.

## METHODS

### Yeast Strains and Plasmids

Standard molecular biology protocols were used to engineer all *S. cerevisiae* strains (Supplementary Table 1) used in this study (Sambrook and Russell 2001). The overall methodology for construction of the triple gene drive strain utilized both standard HR-based chromosomal integrations (*sans* any DSB) and Cas9-based editing (Supplementary Fig. 1). Briefly, DNA constructs were first assembled onto *CEN*-based plasmids (typically pRS315) using *in vivo* assembly in yeast (FINNIGAN AND Thorner 2015). If necessary, point mutations were introduced using PCR mutagenesis (Zheng *et al.* 2004). Next, the engineered cassette was amplified with a high-fidelity polymerase (KOD Hot Start, EMD Millipore), transformed into yeast using a modified lithium acetate method (Eckert-Boulet *et al.* 2012), and integrated at the desired genomic locus. PCR was used to diagnose proper chromosomal position for each integration event followed by DNA sequencing. The DNA maps for manipulated yeast loci are included in Supplementary Fig. 2. DNA plasmids used in this study can be found in Supplementary Table 2. Expression cassettes for sgRNA were based on a previous study (DiCarlo *et al.* 2013), purchased as synthetic genes (Genscript), and sub-cloned to high-copy plasmids using unique flanking restriction sites. All vectors were confirmed by Sanger sequencing.

### Culture Conditions

Budding yeast were cultured in liquid or solid medium. YPD-based medium included 2% peptone, 1% yeast extract, and 2% dextrose. Synthetic (drop-out) medium included yeast nitrogen base, ammonium sulfate, and amino acid supplements. A raffinose/sucrose mixture (2%/0.2%) was used to pre-induce cultures prior to treatment with galactose (2%). All media was autoclaved or filter sterilized (sugars).

### Cas9-based editing *in vivo*

Editing of haploid *S. cerevisiae* strains was performed as previously described (Roggenkamp *et al.* 2018). Briefly, an integrated copy of *S. pyogenes* Cas9 was designed with two flanking “unique” (u2) sites—23 base pairs artificially introduced into the genome. This sequence contains a maximum mismatch to the native yeast genome and is used in order to (i) multiplex at two separate sites using a single guide RNA construct, (ii) minimize (or likely eliminate) potential off-target effects, and (iii) allow for increased biosecurity in testing of active CRISPR gene drive systems (Finnigan and Thorner 2016). Haploid yeast were pre-induced overnight in a raffinose/sucrose mixture to saturation, back-diluted to an OD600 of approximately 0.35 in rich medium containing galactose, and cultured for 4.5 hr at 30°C. Equimolar amounts (1,000 ng) of high-copy plasmid (sgRNA) were transformed into yeast followed by recovery overnight in galactose and a final plating onto SD-LEU medium. Colonies were imaged and quantified after 3-4 days of growth. Haploid yeast editing experiments included three replicates in triplicate—all as separate transformation events—for each strain (n=9).

### Gene drive activation and containment

Haploid yeast strains harboring the gene drive (Cas9) system were first transformed with the sgRNA-containing plasmid (LEU2-marked). Next, drive strains were mated to target strains of the opposite mating type on rich medium for 24 hr. Third, yeast were velvet-transferred to synthetic drop-out medium to select diploids (e.g. SD-URA-LEU or SD-URA-LEU-HIS); each haploid genome contained at least one unique selectable marker. Diploids were selected three consecutive rounds with 1-2 days incubation at each step. Fourth, yeast were cultured in pre-induction medium (raffinose/sucrose) lacking leucine overnight, back-diluted into rich medium containing galactose, and grown for 5 hr (or appropriate time intervals). Strains were diluted to approximately 100-500 cells per mL and plated onto SD-LEU for 2 days. Finally, colonies were transferred to the appropriate selection plates (e.g. SD-HIS, G418, SD-URA, and a fresh SD-LEU plate) for 1 additional day of growth before being imaged. The number of surviving colonies on each media type was quantified; experiments were performed in at least triplicate.

A number of safeguards were included in the design of all gene drive systems. First, the genomic targets for all guide RNAs included only non-yeast sequences (u1, GFP, and Kan^R^) (Finnigan and Thorner 2016; Roggenkamp *et al.* 2017). Second, the primary guide RNA cassette (u1) for targeting of the *HIS3* locus which included Cas9 was maintained on an unstable high-copy (2μ) plasmid; previous work has demonstrated loss of this vector type in the absence of active selection (DiCarlo *et al.* 2015; Roggenkamp *et al.* 2018). Third, the *S. cerevisiae* BY4741/BY4742 genetic background does not readily undergo sporulation, even under optimal conditions. Fourth, Cas9 expression was repressed by growth on dextrose until gene drives were activated. And finally, all diploid strains, plates, and consumables were autoclaved and inactivated.

### Graphics and Evolutionary Analysis

Molecular graphics were generated using the Chimera software package from the Univ. of California, San Francisco (Pettersen *et al.* 2004). Homologous sequences to the yeast DNA Ligase IV (Dnl4) protein were obtained using multiple BLAST (NCBI) searches within either the fungal or metazoan clade (Supplementary Table 3). The phylogenetic tree of DNA Ligase IV was created using the Phylogeny.fr software (DEREEPER *et al.* 2008; DEREEPER *et al.* 2010). Multiple sequence alignments were performed using Clustal Omega (LI *et al.* 2015). The predicted structures of the human, yeast, and mosquito Ligase IV enzyme were generated using I-TASSER (Roy *et al.* 2010). The template structures included the human Lig4 N-terminus (PDB:3W1B) (OCHI *et al.* 2013) and the yeast Dnl4 C-terminus (PDB:1Z56) (DORE *et al.* 2006). Predicted models were individually aligned against the crystal structures using MatchMaker in Chimera. Metrics for the predicted structures are included in Supplementary Table 4.

## ACKNOWLEDEGEMENTS & FUNDING

We thank Cory Wasko and Emily Wedeman (Kansas State University) and Muriel Eaton (Purdue University) for useful comments and Megan Halloran (Kansas State University) for laboratory assistance. This project was supported by an Institutional Development Award (IDeA) from the National Institute of General Medical Sciences of the National Institutes of Health under grant number P20 GM103418 to G.C.F. This work was also supported by the USDA National Institute of Food and Agriculture, Hatch Project 1013520 to G.C.F. The content is solely the responsibility of the authors and does not necessarily represent the official views of the National Institute of General Medical Sciences or the National Institute of Health. Y.Y. and G.C.F built all reagents (plasmids and yeast), performed all experiments, and performed data analyses and figure preparation. G.C.F. wrote the manuscript.

## ANIMAL AND HUMAN SUBJECT STATEMENT

This study does not use any animals or human subjects.

## ABBREVIATIONS

CRISPR: clustered regularly interspaced short palindromic repeats;
NHEJ: non-homologous end joining;
HR: homologous recombination;
DSB: double stranded break;
sgRNA: single guide RNA fragment;
GD: CRISPR-based gene drive system;
CGM: “complete” multi-locus drive;
MGD: “minimal” multi-locus drive;
BRCT: BRCA1 C-Terminal domains;
UTR: untranslated region;
indel: insertion or deletion.

